# De novo mutations drive the spread of macrolide-resistant *Mycoplasma genitalium*: a mathematical modelling study

**DOI:** 10.1101/321216

**Authors:** Dominique Cadosch, Victor Garcia, Christian L. Althaus, Jørgen Skov Jensen, Nicola Low

## Abstract

**Background:** The rapid spread of azithromycin resistance in sexually transmitted *Mycoplasma genitalium* infections is a growing concern. It is not yet clear to what degree macrolide resistance in *M. genitalium* results from the emergence of de novo mutations or the transmission of resistant strains.

**Methods:** We analyzed epidemiological data and developed a compartmental model to investigate the contribution of de novo macrolide resistance mutations to the spread of antimicrobial-resistant *M. genitalium*. We fitted the model to data from France, Denmark and Sweden and estimated treatment rates of infected individuals and the time point of azithromycin introduction.

**Results:** We found a high probability of de novo resistance (12%, 95% CI 8–17%), which is responsible for the observed rapid spread of antimicrobial resistant *M. genitalium*. The estimated per capita treatment rate in France was lower than in Denmark and Sweden but confidence intervals for the three estimates overlap. The estimated dates of introduction of azithromycin in each country are consistent with published reports.

**Conclusions:** Since de novo resistance is the main driver of macrolide resistance in *M. genitalium*, blind treatment of urethritis with azithromycin is not recommended. Clinical management strategies for *M. genitalium* should limit the unnecessary use of macrolides.

## Introduction

Macrolide-resistant *Mycoplasma genitalium* already accounts for 40% or more of detected infections in some countries [1–4]. *M. genitalium* is a sexually transmitted bacterium which, together with *Chlamydia trachomatis*, causes non-gonococcal urethritis (NGU) in men [5] and lower and upper genital tract disease in women [6]. *M. genitalium* is detected using nucleic acid amplification tests (NAATs) [7], which were first developed during the 1990s as research tools because the bacterium is slow-growing and hard to culture. In most clinical settings, NAATs for *M. genitalium* diagnosis are not available. The clinical syndrome of NGU is treated empirically, with a single 1g dose of azithromycin, recommended for first line treatment in many countries since the late 1990s [8].

Macrolide resistance in *M. genitalium* results from a single nucleotide mutation in region V of the 23S rRNA gene, most commonly A2058G or A2059G. Jensen et al. identified these mutations in Australian and Swedish men, with NGU caused by *M. genitalium*, who experienced clinical treatment failure with 1g azithromycin [9]. The men carried a wild-type organism before treatment, but post-treatment specimens contained mutations in the 23S rRNA gene that conferred macrolide resistance [9]. Since then, other investigators have detected macrolide resistance mutations de novo (also known as acquired, induced or selected) in *M. genitalium* [10–15]. Once acquired, untreated resistant strains can be transmitted to new sexual partners.

Recommendations for future research on *M. genitalium* prioritize the need for more effective and safe antimicrobials [16]. It is important to understand the degree to which treatment failure in *M. genitalium* results from the emergence of de novo resistance mutations or the transmission of resistant strains because the type of resistance will influence future strategies. The objective of this study was to investigate the role of de novo emergence of resistance in the spread of azithromycin-resistant *M. genitalium*.

## Methods

We developed a mathematical model of *M. genitalium* transmission and fitted it to epidemiological data about time trends in macrolide resistance. We define ‘de novo’ as a change from a drug-susceptible infection before treatment to a drug-resistant infection after treatment, either by selection of one or a few pre-existing resistant mutants in an otherwise drug-susceptible bacterial population or due to a novel resistance mutation evolving during drug exposure. We used *R 3.3.2* for statistical analyses, transmission model simulations and parameter inference.

### Epidemiological data

We searched Pubmed in March 2018 and updated the search on 4^th^ May 2018. We used the medical subject headings *Mycoplasma genitalium* AND *drug resistance, bacterial* and found 67 publications. Two authors independently searched for countries with multiple studies that reported on *M. genitalium* and macrolide resistance mutations. We selected three countries with data for more than three years from the same region or an entire country, and which used different strategies to test and treat *M. genitalium*. For each country, we recorded the testing strategy and treatment regimen, year in which azithromycin was introduced for *M. genitalium* treatment, numbers of specimens with positive results for *M. genitalium* and the number with macrolide resistance mutations. We contacted study authors for additional information. For each year, we calculated the proportion (with 95% CI) of azithromycin-resistant *M. genitalium*.

### Mathematical model

We developed a mathematical model that simulates the spread of drug resistance within a population (Figure 1). The model consists of three compartments: susceptibles (S), people infected with a drug-susceptible strain of *M. genitalium* (I_S_), and people infected with a drug-resistant strain of *M. genitalium* (I_R_).

**Figure 1.**
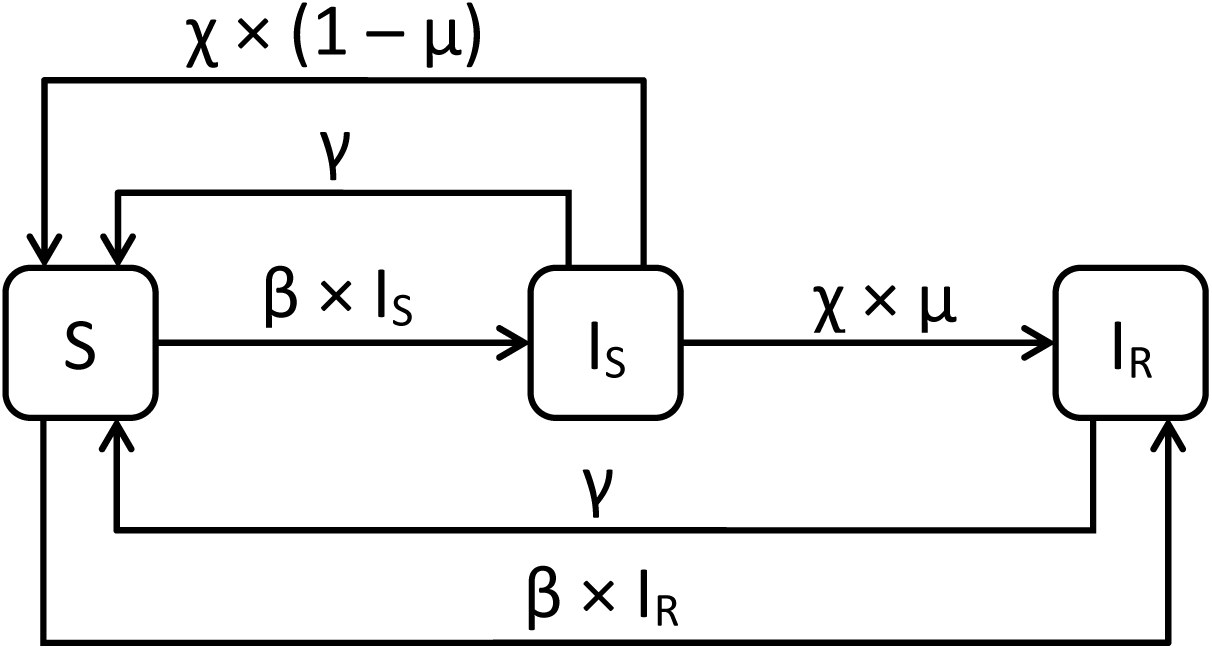
Structure of the epidemiological model for *M. genitalium*.

Assuming a homogenous population without demography, the transmission dynamics can be described by the following ordinary differential equations:

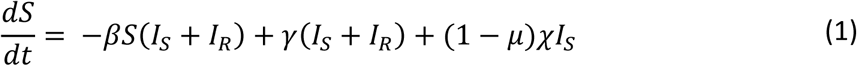

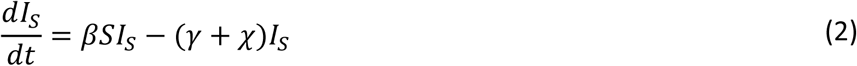

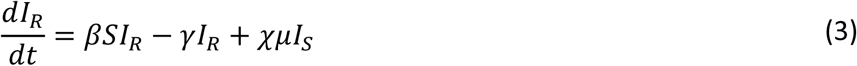

where *β* is the transmission rate, which is assumed to be independent of the type of *M. genitalium* strain. Both types of infections can clear naturally at a rate *γ*. Patients receive treatment at rate *χ*. The treatment rate is defined as all occasions of treatment with a single 1g dose of azithromycin in a person infected with *M. genitalium*, either with or without symptoms. *µ* denotes the probability of de novo resistance emergence during treatment. The de novo emergence of resistance also implies that the treatment failed. We used the point estimate of the probability of de novo resistance emergence from a meta-analysis, described below. For simplicity, we assumed that resistant infections clear naturally, with no second-line treatment. The rate at which the drug-resistant strain replaces the drug-susceptible in a population can be expressed by the difference in the net growth rates (Δ*φ*) between the two strains [27, 28]:

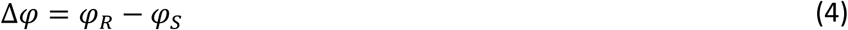

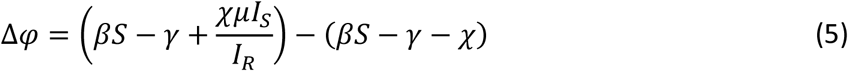

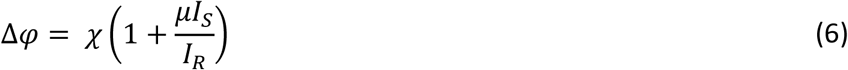

### Model parameters

We set the natural clearance rate (*γ*) of *M. genitalium* to 0.8 y^-1^ [19], and the infection rate, *β,* to 0.816 person^-1^ *y*^*-1*^. These values result in an equilibrium prevalence of *M. genitalium* infections of about 2% in the absence of treatment, which is consistent with estimates of the prevalence of *M. genitalium* in sexually active adults in high-income countries [20]. The values for the natural clearance rate and the prevalence of infection do not govern the relative growth rate of the drug-resistant proportion (Equation 6), so they do not influence the relative prevalence of resistant infections or estimates of the treatment rate. We did not find any published evidence of the effect of macrolide resistance on the fitness of *M. genitalium* strains, so we assumed that any fitness reduction is negligible and that resistant and wild-type strains have the same infectivity.

The probability of emergence of de novo resistance during treatment, *μ*, was derived from a meta-analysis of estimates from published studies (supplementary table 1). Our search identified one study [15] published since the search limit of a previous meta-analysis [21]. All included studies investigated patients with *M. genitalium* who received a single 1g dose of azithromycin and who had both pre-and post-treatment specimens tested for macrolide resistance mutations. From each study we extracted the total number of patients with a wild-type pretreatment sample and the number with post-treatment resistance. In contrast to Horner et al. [21], we excluded three patients from one study [13] because we could not confirm the inclusion criteria. We applied the Freeman-Tukey double arcsine transformation to the calculated proportions [22,23] and used a random effects model to estimate the average proportion (with 95% confidence intervals, CI) of patients with initially macrolide susceptible *M. genitalium* who had macrolide resistance mutations detected after treatment (*metaprop* function from the *R* package *meta 4.9*). We estimated that de novo resistance develops in 12% (95% CI 8–17%, I^2^ 34%) of *M. genitalium* infections treated with 1g azithromycin.

**Figure 2.**
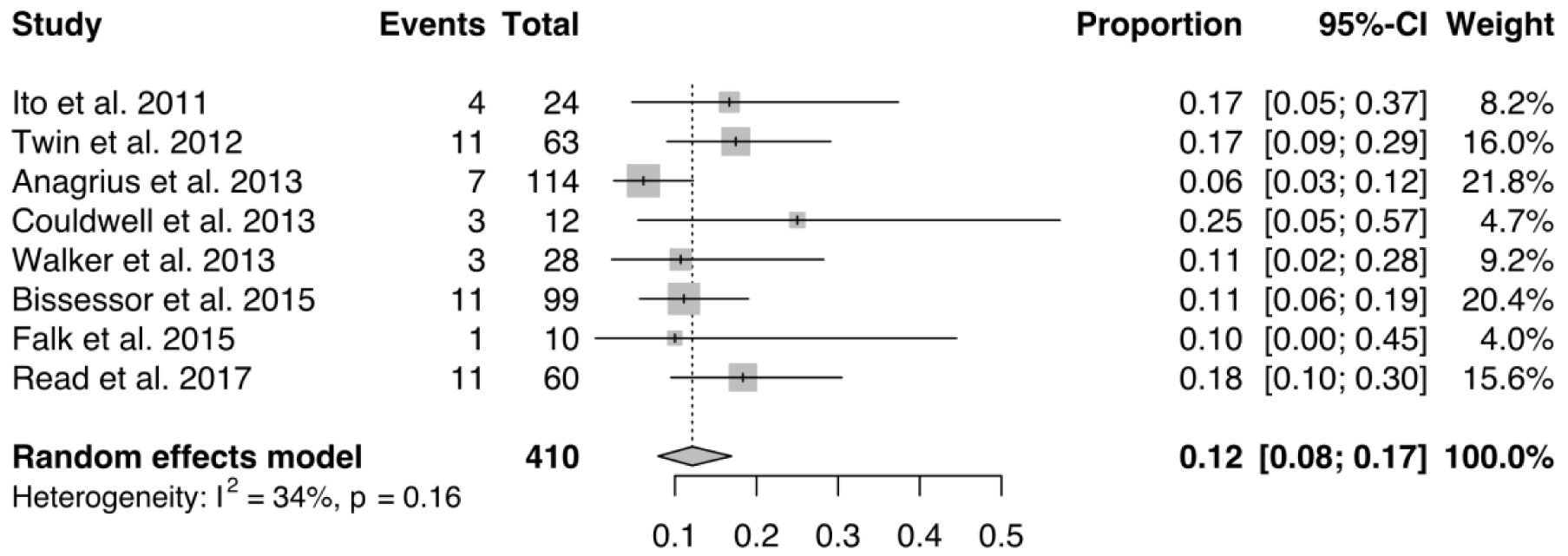
Probability of de novo emergence of azithromycin resistance in *M. genitalium*, estimated by random effects meta-analysis of treatment studies reporting pre-treatment susceptibility to azithromycin [10,11,13–15,21].

**Table 1.** Model parameters

| Parameter | Description | Value | Source |
| --- | --- | --- | --- |
| $\beta$ | Transmission rate | 0.816 person<sup>-1</sup> y<sup>-1</sup> | See text |
| $\gamma$ | Natural clearance rate | 0.8 y<sup>-1</sup> | [19] |
| $\chi$ | Treatment rate of infecteds | Estimated | |
| $\mu$ | Probability of de novo resistance during treatment | 12%<br>(95% CI: 8 – 17%) | Meta-analysis of<br>[10–15,21], see<br>Figure 2 |

### Model fitting and simulations

We fitted the transmission model to country-specific resistance data to obtain maximum likelihood estimates of the treatment rate of infected people *χ* and the time point *T* for the introduction of azithromycin. Given a model-predicted proportion of resistant strains 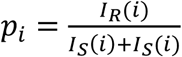 in year *i*, the log-likelihood to find *k*_*i*_ resistant samples in *N*_*i*_ tested individuals is:

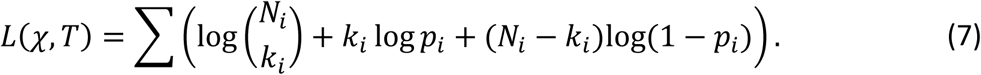

Simulations start at time *T* with 98% uninfected people, 2% people with drug susceptible infections and no people with initial drug-resistant infections. We used log-transformed parameters for the estimation and stipulated that the upper limit of *T* could not be beyond the time point where resistance was first observed. We derived simulation-based 95% confidence intervals for the model curve from 10,000 bootstrap samples from the multivariate normal distribution of the two parameters. We used the *ode* function from the *R* package *deSolve 1.20* to solve the ordinary differential equations, and the *mle2* function from the *R* package *bbmle 1.0.19* using the *Nelder-Mead* method for log-likelihood optimization.

To investigate the influence of the level of de novo resistance emergence on the rise in the proportion of resistant infections, we simulated two alternative scenarios. In these scenarios, we kept the model-derived maximum likelihood estimates of χ and *T* but set the probability of de novo resistance emergence to lower values (*μ* = 1% and *μ* = 0.1%). As a sensitivity analysis we fitted the model to the country-specific data while assuming lower de novo resistance emergence probabilities of *μ* = 1% and *μ* = 0.1% and estimated both the resulting alternative χ and *T* values and the goodness of fit. The goodness of fit was measured with the aggregated negative log-likelihoods of the model fits relative to the country-specific data and the Akaike information criterion (AIC) [24].

## Results

### Data

We included five studies that provided data about the proportion of azithromycin-resistant *M. genitalium* infections over time and the management of *M. genitalium* infection in France [25–27], Denmark [1] and Sweden [11] (supplementary table 2). Study authors provided additional information from Denmark (data disaggregated by year) and Sweden (numbers of patients per year and unpublished data for 2012 and 2013) [1,11].

In France, we included three studies with data from 314 patients (310 from Bordeaux) from 2003 to 2012 [25–27]. None of 17 *M. genitalium* positive specimens from 2003 to 2005 contained macrolide resistance mutations. From 2006 onwards, mutations were detected in 10% to 17% of specimens tested in each year. In France, azithromycin was introduced for first line treatment of NGU in the 1990s [28]. For Denmark, one study reported nationwide data from 1,008 patients with *M. genitalium* detected from 2006 to 2010, with 27% to 42% of specimens containing macrolide resistance mutations [1]. In Denmark, 1g single dose azithromycin is routinely prescribed for treatment of NGU; erythromycin was the first-line treatment before azithromycin became available. An extended azithromycin regimen is prescribed if a *M. genitalium* infection was diagnosed and NAAT for detection of *M. genitalium* infections have been available since 2003 [1]. In Sweden, we analyzed one study with data about macrolide resistance mutations from 408 samples obtained from 2006 to 2013 from patients at a single clinic in Falun [11]. Macrolide resistance mutations were first detected in a single specimen in 2008 and increased to 16% of 95 specimens in 2011. In Sweden, doxycycline is used as first line treatment for NGU [29]. Azithromycin is used only when *M. genitalium* is identified as the cause, with testing introduced in the 2000s [11].

### Mathematical model

The transmission model fitted the increase in *M. genitalium* resistance in France, Denmark and Sweden well (Figure 3). The rise in the proportion of azithromycin resistant *M. genitalium* infections in all three countries was consistent with de novo emergence of macrolide resistance mutations in 12% of initially wild-type infections. The alternative scenarios show that lower probabilities of de novo resistance, but the same estimated treatment rates and time points for the introduction of azithromycin, would result in considerably lower proportions of resistant *M. genitalium* infections (Figure 3).

**Figure 3.**
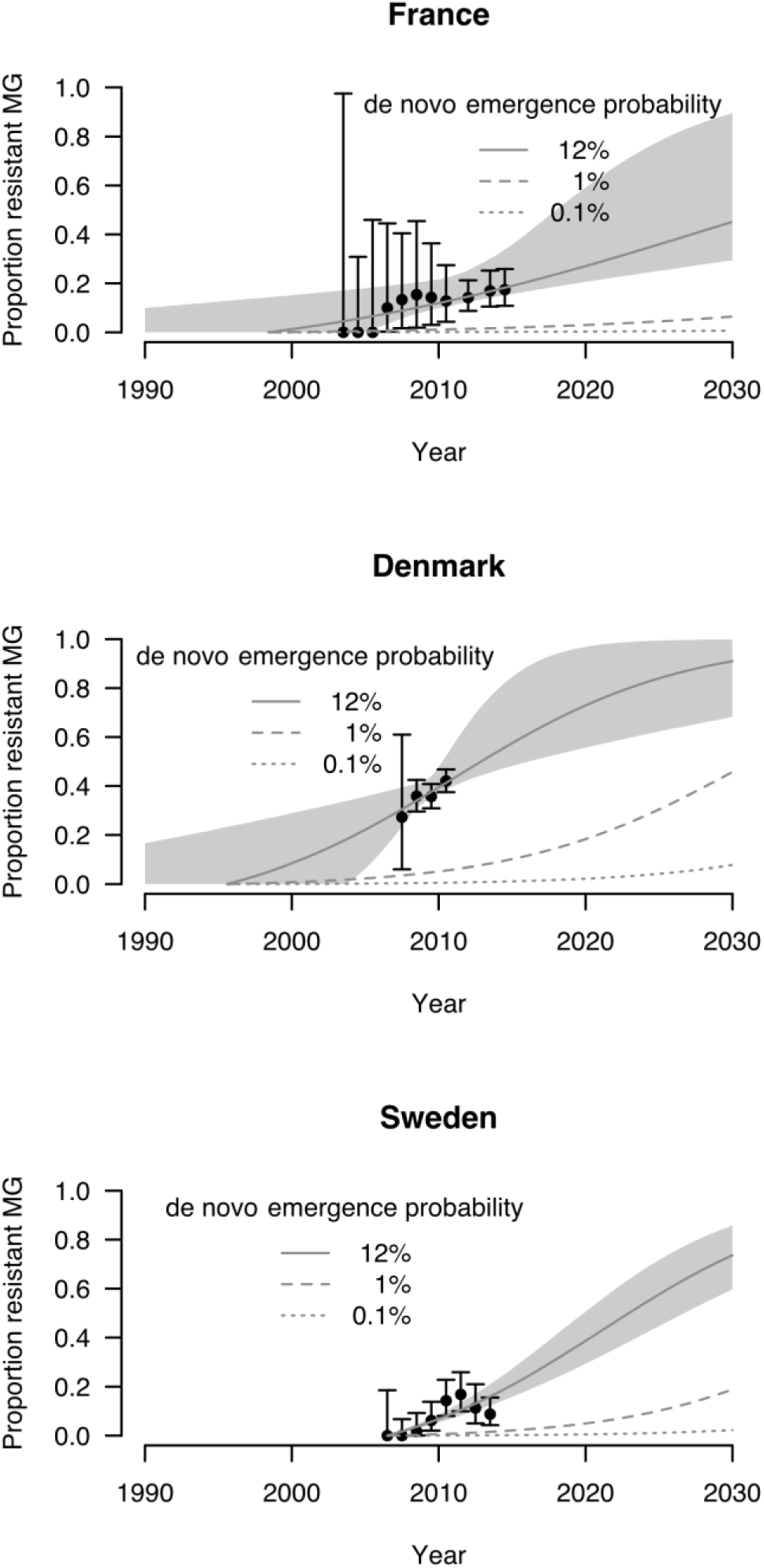
Maximum likelihood fits (solid grey lines) of the transmission model to the data of the relative prevalence of azithromycin-resistant *M. genitalium* infections in France, Denmark and Sweden over time. The black data points correspond to reported proportions of resistant infections [1,11,25–27] (including unpublished data from Denmark and Sweden). The error bars indicate the 95% confidence intervals. The light grey area is the 95% confidence interval of the model predictions. The dashed and dotted lines depict alternative scenarios with lower probabilities of de novo emergence of resistance but with the treatment rate and time point of azithromycin introduction obtained from the maximum likelihood fit.

The model estimated treatment rate of infected people and date of introduction of azithromycin were: France, treatment rate 0.07 y^-1^ (95% CI 0.02 – 0.18 y^-1^),introduction of azithromycin in May 2000 (95% CI October 1986 – June 2005); Denmark, treatment rate of 0.13 y^-1^ (95% CI 0.05 – 0.34 y^-1^), introduction of azithromycin in August 1996 (95% CI November 1976 – January 2004); Sweden, treatment rate 0.14 y^-1^ (95% CI 0.11 – 0.17 y^-1^), introduction of azithromycin July 2006 (95% CI January 2006 – November 2006). A treatment rate of 0.14 y^-1^, such as in Sweden, corresponds to a proportion of 1 – *e*^-0.14^ ∼ 13% of infected individuals that will have received treatment after one year.

In the sensitivity analysis, assuming de novo resistance emergence probabilities of 1% or 0.1%, the model estimated higher treatment rates of infected individuals, implausibly early time points of azithromycin introduction and worse goodness of fit (supplementary table 3).

The probability of de novo resistance emergence increases the growth rate of resistant infections across different initial values for the proportion of resistant infections (Figure 4). The relationship between the proportion of drug-resistant *M. genitalium* infections (Δ*φ*) and the proportion of resistant infections explains some of the dynamics of resistance spread. The growth advantage conferred by de novo emergence of resistant strains is always greatest at the time of introduction of antibiotic treatment, when the proportion of resistant strains is the lowest. As the resistant strain spreads, the growth advantage diminishes, slowly approaching Δ*φ* = χ according to Equation 6. Thus, the growth acceleration of resistant strains provided by de novo resistance reduces as the resistant strain spreads through the pathogen population. The curves for which we assumed a lower de novo resistance emergence probability are substantially flatter than the curve that results from the probability of de novo resistance estimated from the data.

**Figure 4.**
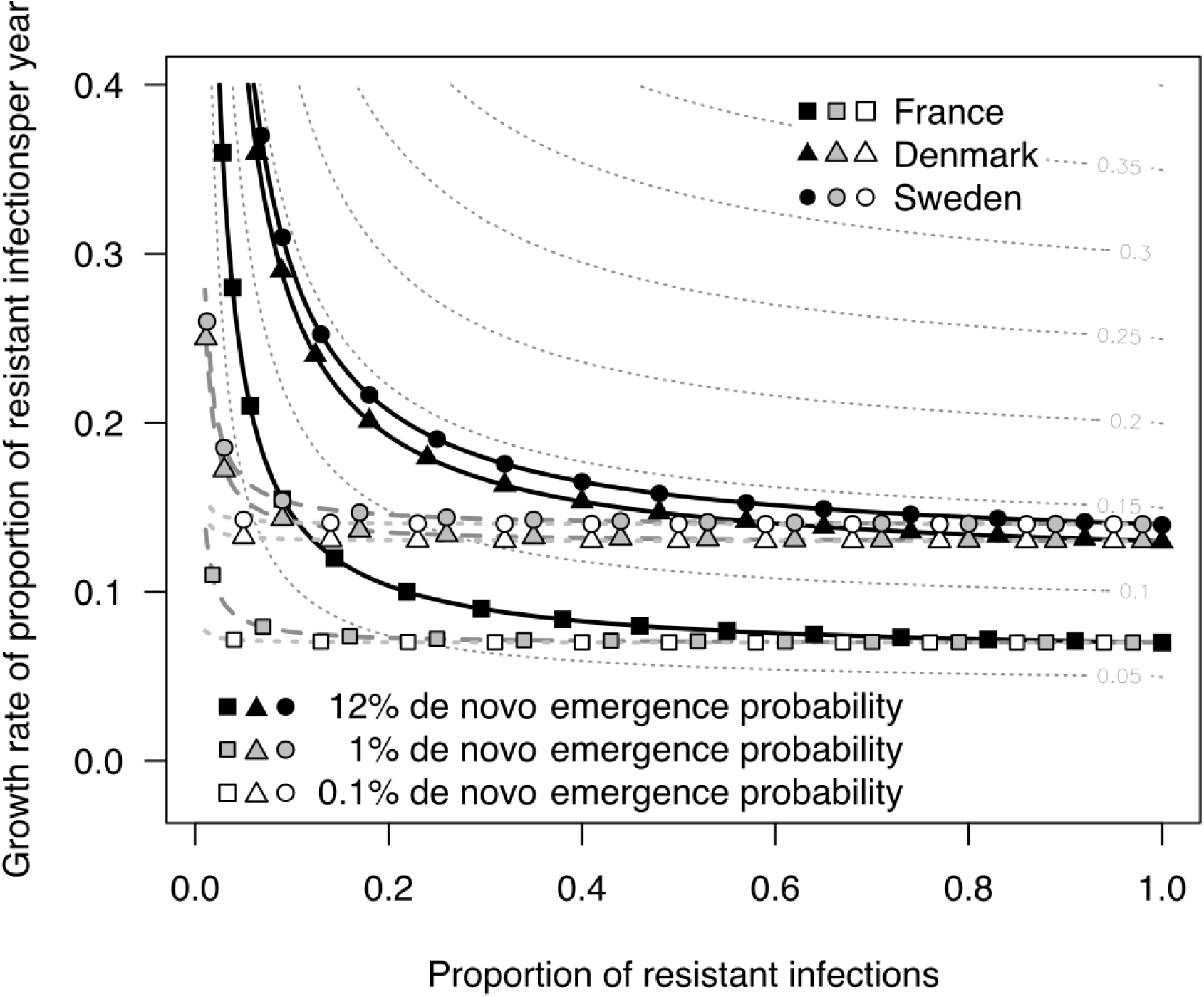
Relationship between the growth rate of the drug-resistant proportion of *M. genitalium* infections and the proportion of resistant infections for different treatment rates. The colored solid lines show the growth rate for France (squares), Denmark (triangles), and Sweden (circles) in a model that includes the established 12% probability for de novo emergence of drug resistance (black symbols). The grey and white symbols are the growth rates in a model that uses the same treatment rate but reduced probabilities for de novo emergence of resistance.

## Discussion

In this study, we fitted a compartmental model to time trend data about the proportions of azithromycin-resistant *M. genitalium* infections in France, Denmark and Sweden and estimated the treatment rates and the time point of introduction of azithromycin in those countries. Alternative scenarios with the same model would result in considerably slower rise of *M. genitalium* resistance.

This study strongly suggests that, rather than resulting in ‘occasional treatment failure’ as originally believed [9], the development of de novo resistant mutations in about one in eight *M. genitalium* infections [21] is the main driver of azithromycin resistance. The data from France and Sweden [11,25–27], where no macrolide resistant mutations were detected initially, show a substantial proportion of diagnosed *M. genitalium* infections with azithromycin resistance after just a few years of azithromycin use. Our model shows that a high de novo resistance acquisition rate contributes considerably to the spread of resistance, particularly during the early stages. The effect then decreases as the proportion of resistant infections increases. This pattern contrasts with alternative scenarios in which resistance emerges with a lower probability. Then, the effect on the growth rate would be substantially smaller and the growth dynamics of the drug resistant proportion are much closer to a logistic growth model. Assuming the same treatment rate of infected individuals and time point of introduction of azithromycin, this growth dynamic would require much more time to reach the levels of resistance that we observed. Our model-predicted estimates of the introduction of azithromycin for the treatment of NGU were consistent with published data describing its use in France [28] and Denmark in the 1990s, but later introduction in Sweden [11]. Our estimated treatment rate of infected individuals for France was lower than those for Denmark and Sweden but the 95% confidence intervals of all three estimates overlap. The estimated rates in Sweden and Denmark are very close to those estimated in another epidemiological model of *M. genitalium* infections in the United Kingdom [30].

The high probability of de novo emergence of macrolide resistance mutations during treatment of *M. genitalium* infections appears to differ from experiences with some other sexually transmitted bacterial infections. A 1g dose of azithromycin might often be insufficient to eradicate a *M. genitalium* infection in concert with host immune responses, allowing for either a resistance mutation to occur in the single 23S rRNA operon during treatment or the survival of a few pre-existing drug-resistant bacteria and the subsequent selection of the mutants. The latter explanation is favored by the strong association with de novo resistance and high organism load [10,15], but both mechanisms may play a role. In the absence of any observable fitness cost, or of routine tests to detect macrolide resistance mutations, *M. genitalium* resistance has emerged and spread rapidly. In contrast, selection pressure exerted by treatment and clonal spread are the major drivers of the spread of macrolide-resistant *Neisseria gonorrhoeae,* with de novo resistance considered to be negligible [18]. *N. gonorrhoeae* has four copies of the 23S rRNA gene and resistance increases with the number of mutated copies [31]. In addition, active measures are used to limit the potential for the emergence of de novo macrolide resistance in *N. gonorrhoeae*, including dual therapy, in which azithromycin is a second drug in combination with ceftriaxone. Transmitted resistance is assumed to be responsible for most antimicrobial resistance, but a high rate of de novo resistance emergence has been observed during treatment with various antibiotics of infections such as *Pseudomonas aeruginosa* and *Enterobacteriaceae* [32,33]. In general, de novo selection of drug-resistant mutants within a single patient occurs more often if the resistance is mediated by single-base mutations than if acquisition of efflux pumps or other complex mechanism are needed [34]. Thus, it is distinct from the selection of drug resistance as a result of treatment at the population level which is more often transmitted; a situation which is seen with most other bacterial and parasitic sexually transmitted infections.

A strength of this study is the use of empirical data sources and mathematical modelling. Parameters that were not available in the literature were estimated by fitting the model to observational data. Despite its simplicity, the model assumptions provide a coherent qualitative explanation for the clinically observed rapid rise of macrolide resistant *M. genitalium* infections.

There are some caveats to both the observational data sources and the model. First, owing to the small number of samples for each data point, particularly for early years, confidence intervals for those estimates of the proportion of resistant infections are wide. In Denmark, azithromycin has been used for a long time but data about the prevalence of drug resistant infections were only available since 2006, which introduces more uncertainty in the estimated point at which resistance emerged. Second, the characteristics of people tested for *M. genitalium* in the three countries are not well described and differences in testing practices between countries might account for some of the variation in the proportions with macrolide resistance. An increase over time in the proportion of resistant infections was, however, observed in all three countries. We used a relatively simple transmission model, so we made several simplifying assumptions. First, we assumed that treatment rates of infected individuals in each country were constant over time and did not account for the possibility that azithromycin use might have changed over time. Second, we assumed that no second line treatments were used for resistant *M. genitalium* infections. In practice, since most *M. genitalium* infections are asymptomatic and diagnostic testing is still uncommon, we do not think that this simplification affected our results. Third, our model does not include detailed population structure because the rate at which the relative proportion of resistant bacterial strains spread in a population can often be explained by the treatment rate, rather than the sexual network structure [18]. More complex models with different sexes, partner change rates and age structure, would be necessary to obtain a better description of the absolute prevalence of infections and resistance, but this was not the objective of this study.

The high level of azithromycin resistance in *M. genitalium*, driven by de novo resistance, poses problems for clinical management and population level control strategies [35]. Screening and treatment of asymptomatic *M. genitalium* would simply result in the spread of more strains with de novo mutations, with absent evidence of a reduction in clinical morbidity [35]. Treatment strategies to maintain the use of existing antimicrobials are also being evaluated since resistance to second line treatment with moxifloxacin is already increasing [3]. In an observational study, resistance-guided therapy for symptomatic *M. genitalium*, with initial treatment with doxycycline followed by 2.5g azithromycin over three days for macrolide susceptible infections and sitafloxacin for resistant infections resulted in an incidence of de novo macrolide resistance of 2.6% (95% CI 0.3–9.2%) [36].

Randomized controlled trials are now needed to evaluate different treatment algorithms and new antimicrobials or combination therapy that might have a lower propensity for the emergence of de novo resistance [8]. Since de novo resistance is the main driver of azithromycin resistance in *M. genitalium*, blind treatment of urethritis with single dose azithromycin is not recommended. Clinical management strategies for *M. genitalium* and other STIs should seek to limit the unnecessary use of macrolides.

## Acknowledgement

We would like to thank Carin Anagrius from the Falu lasarett in Falun, Sweden and Kirsten Salado-Rasmussen from the Bispebjerg Hospital in Copenhagen, Denmark for providing us with additional unpublished data.

## Previous presentation

An earlier version of this work was presented as a poster at EPIDEMICS 6: 6th International Conference on Infectious Disease Dynamics. Sitges, Spain, November 29 to December 1 2017. Cadosch D, Garcia V, Althaus CL, Low N. Inadequate treatment accelerates the spread of macrolide resistance in Mycoplasma genitalium

## Supplementary material

**Supplementary table S1.**
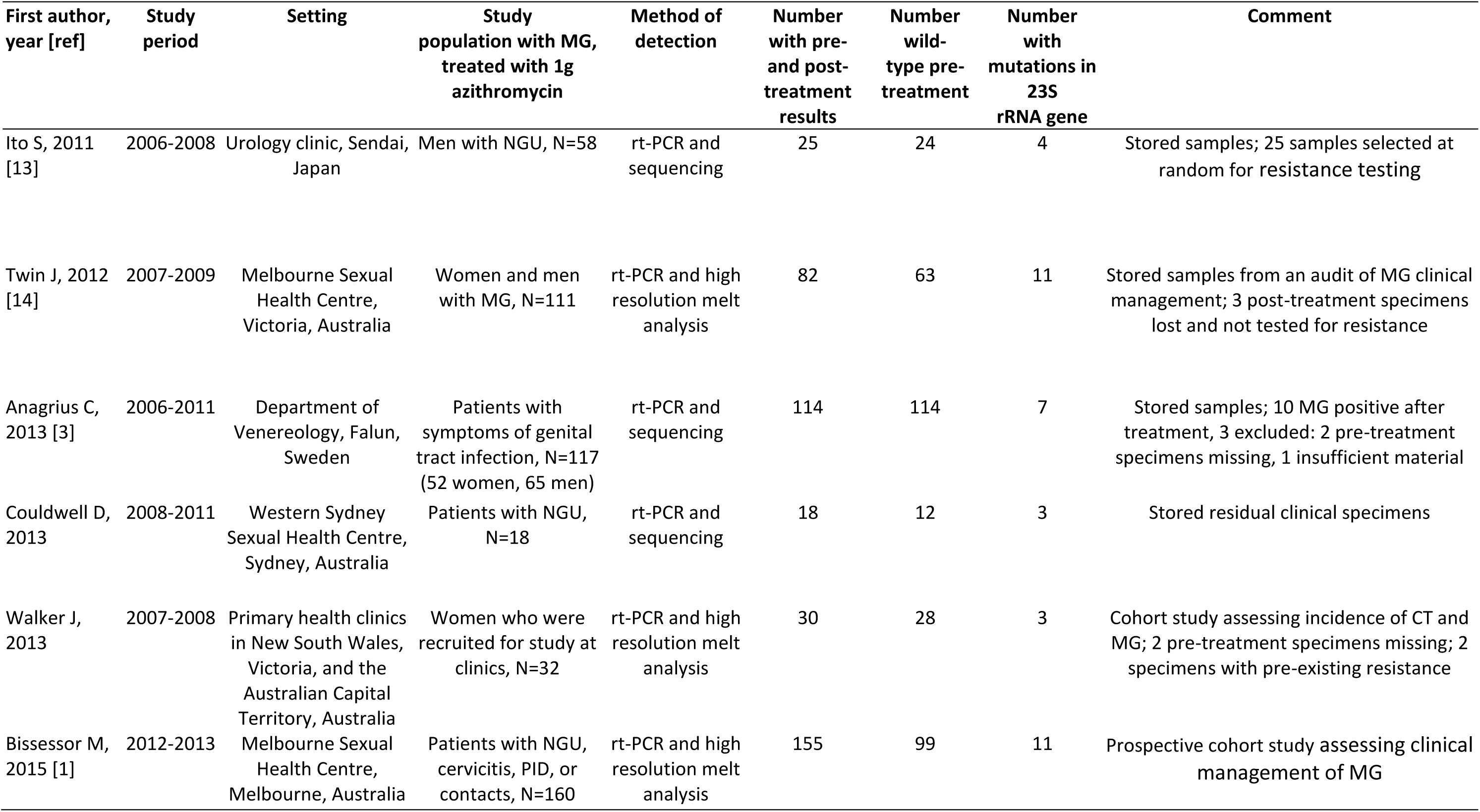
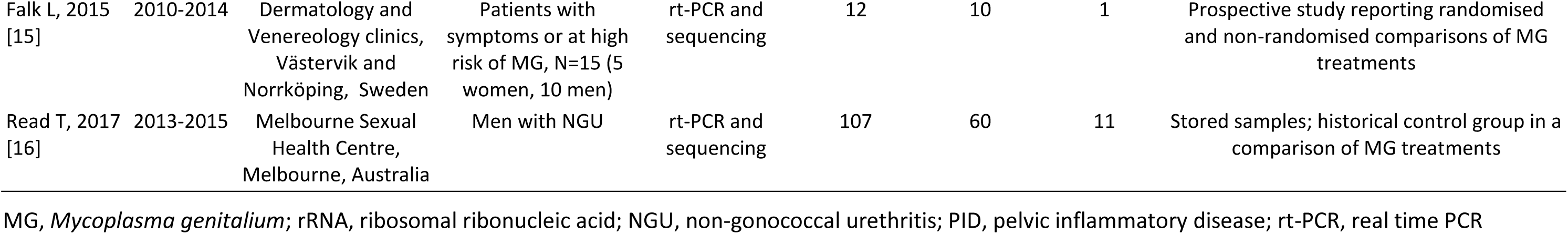
Characteristics of studies included in meta-analysis, by year of publication

| First author, year [ref] | Study period | Setting | Study population with MG, treated with 1g azithromycin | Method of detection | Number with pre- and post-treatment results | Number wild-type pre-treatment | Number with mutations in 23S rRNA gene | Comment |
| --- | --- | --- | --- | --- | --- | --- | --- | --- |
| Ito S, 2011 [13] | 2006-2008 | Urology clinic, Sendai, Japan | Men with NGU, N=58 | rt-PCR and sequencing | 25 | 24 | 4 | Stored samples; 25 samples selected at random for resistance testing |
| Twin J, 2012 [14] | 2007-2009 | Melbourne Sexual Health Centre, Victoria, Australia | Women and men with MG, N=111 | rt-PCR and high resolution melt analysis | 82 | 63 | 11 | Stored samples from an audit of MG clinical management; 3 post-treatment specimens lost and not tested for resistance |
| Anagrus C, 2013 [3] | 2006-2011 | Department of Venereology, Falun, Sweden | Patients with symptoms of genital tract infection, N=117 (52 women, 65 men) | rt-PCR and sequencing | 114 | 114 | 7 | Stored samples; 10 MG positive after treatment, 3 excluded: 2 pre-treatment specimens missing, 1 insufficient material |
| Couldwell D, 2013 | 2008-2011 | Western Sydney Sexual Health Centre, Sydney, Australia | Patients with NGU, N=18 | rt-PCR and sequencing | 18 | 12 | 3 | Stored residual clinical specimens |
| Walker J, 2013 | 2007-2008 | Primary health clinics in New South Wales, Victoria, and the Australian Capital Territory, Australia | Women who were recruited for study at clinics, N=32 | rt-PCR and high resolution melt analysis | 30 | 28 | 3 | Cohort study assessing incidence of CT and MG; 2 pre-treatment specimens missing; 2 specimens with pre-existing resistance |
| Bissessor M, 2015 [1] | 2012-2013 | Melbourne Sexual Health Centre, Melbourne, Australia | Patients with NGU, cervicitis, PID, or contacts, N=160 | rt-PCR and high resolution melt analysis | 155 | 99 | 11 | Prospective cohort study assessing clinical management of MG |
| Falk L, 2015<br>[15] | 2010-2014 | Dermatology and<br>Venereology clinics,<br>Västervik and<br>Norrköping, Sweden | Patients with<br>symptoms or at high<br>risk of MG, N=15 (5<br>women, 10 men) | rt-PCR and<br>sequencing | 12 | 10 | 1 | Prospective study reporting randomised<br>and non-randomised comparisons of MG<br>treatments |
| Read T, 2017<br>[16] | 2013-2015 | Melbourne Sexual<br>Health Centre,<br>Melbourne, Australia | Men with NGU | rt-PCR and<br>sequencing | 107 | 60 | 11 | Stored samples; historical control group in a<br>comparison of MG treatments |

MG, *Mycoplasma genitalium*; rRNA, ribosomal ribonucleic acid; NGU, non-gonococcal urethritis; PID, pelvic inflammatory disease; rt-PCR, real time PCR

**Supplementary table S2.**
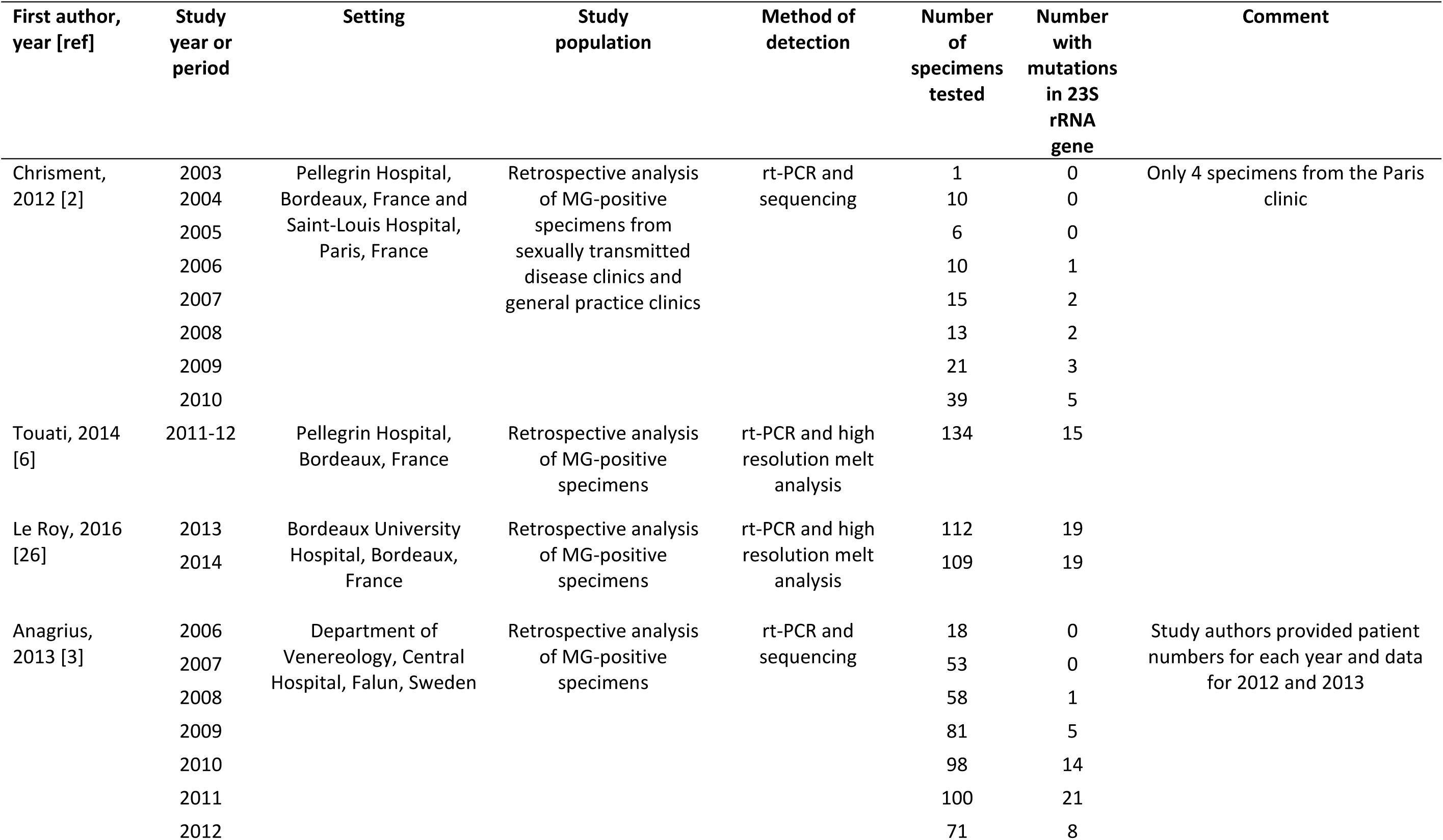
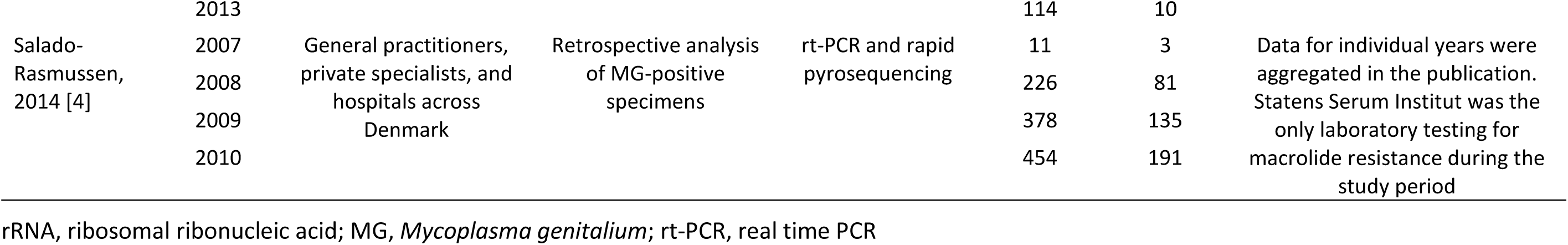
Characteristics of studies with time trend data about azithromycin-resistant infections

| First author, year [ref] | Study year or period | Setting | Study population | Method of detection | Number of specimens tested | Number with mutations in 23S rRNA gene | Comment |
| --- | --- | --- | --- | --- | --- | --- | --- |
| Chrisment, 2012 [2] | 2003 | Pellegrin Hospital, Bordeaux, France and Saint-Louis Hospital, Paris, France | Retrospective analysis of MG-positive specimens from sexually transmitted disease clinics and general practice clinics | rt-PCR and sequencing | 1 | 0 | Only 4 specimens from the Paris clinic |
|  | 2004 |  |  |  | 10 | 0 |  |
|  | 2005 |  |  |  | 6 | 0 |  |
|  | 2006 |  |  |  | 10 | 1 |  |
|  | 2007 |  |  |  | 15 | 2 |  |
|  | 2008 |  |  |  | 13 | 2 |  |
|  | 2009 |  |  |  | 21 | 3 |  |
|  | 2010 |  |  |  | 39 | 5 |  |
| Touati, 2014 [6] | 2011-12 | Pellegrin Hospital, Bordeaux, France | Retrospective analysis of MG-positive specimens | rt-PCR and high resolution melt analysis | 134 | 15 |  |
| Le Roy, 2016 [26] | 2013 | Bordeaux University Hospital, Bordeaux, France | Retrospective analysis of MG-positive specimens | rt-PCR and high resolution melt analysis | 112 | 19 |  |
|  | 2014 |  |  |  | 109 | 19 |  |
| Anagrus, 2013 [3] | 2006 | Department of Venereology, Central Hospital, Falun, Sweden | Retrospective analysis of MG-positive specimens | rt-PCR and sequencing | 18 | 0 | Study authors provided patient numbers for each year and data for 2012 and 2013 |
|  | 2007 |  |  |  | 53 | 0 |  |
|  | 2008 |  |  |  | 58 | 1 |  |
|  | 2009 |  |  |  | 81 | 5 |  |
|  | 2010 |  |  |  | 98 | 14 |  |
|  | 2011 |  |  |  | 100 | 21 |  |
|  | 2012 |  |  |  | 71 | 8 |  |
|  | 2013 |  |  |  | 114 | 10 |  |
| Salado-Rasmussen, 2014 [4] | 2007 | General practitioners, private specialists, and hospitals across Denmark | Retrospective analysis of MG-positive specimens | rt-PCR and rapid pyrosequencing | 11 | 3 | Data for individual years were aggregated in the publication. Statens Serum Institut was the only laboratory testing for macrolide resistance during the study period |
|  | 2008 |  |  |  | 226 | 81 |  |
|  | 2009 |  |  |  | 378 | 135 |  |
|  | 2010 |  |  |  | 454 | 191 |  |

rRNA, ribosomal ribonucleic acid; MG, *Mycoplasma genitalium*; rt-PCR, real time PCR

**Supplementary table S3.** Log-likelihoods of model fits, estimated treatment rates and estimated time points of azithromycin introduction with different probabilities of emergence of *de novo* azithromycin resistance

|  | Probability of emergence of *de novo* azithromycin resistance |  |  |
| --- | --- | --- | --- |
|  | 12% | 1% | 0.1% |
| **France** |  |  |  |
| Log-likelihood | -14.95 | -15.07 | -15.09 |
| Treatment rate person<sup>-1</sup> year<sup>-1</sup> | 0.07 (95% CI 0.02 - 0.18) | 0.10 (95% CI 0.03 - 0.30) | 0.11 (95% CI 0.01 - 1.37) |
| Azithromycin introduction | May 2000<br>(95% CI: Oct 1986 - Jun 2005) | Nov 1985<br>(95% CI: Oct 1941 - Nov 2000) | Apr 1966<br>(95% CI Sep 1473 - Mar 2005) |
| **Denmark** |  |  |  |
| Log-likelihood | -11.21 | -11.20 | -11.20 |
| Treatment rate person<sup>-1</sup> year<sup>-1</sup> | 0.13 (95% CI 0.05 - 0.34) | 0.15 (95% CI 0.07 - 0.32) | 0.16 (95% CI 0.10 - 0.24) |
| Azithromycin introduction | Aug 1996<br>(95% CI Nov 1976 - Jan 2004) | Aug 1983<br>(95% CI Jun 1957 - Jun 1996) | May 1969<br>(95% CI Nov 1947 - Apr 1983) |
| **Sweden** |  |  |  |
| Log-likelihood | -19.00 | -20.86 | -21.07 |
| Treatment rate person<sup>-1</sup> year<sup>-1</sup> | 0.14 (95% CI 0.11 - 0.17) | 0.21 (95% CI 0.11 - 0.40) | 0.22 (95% CI 0.11 - 0.43) |
| Azithromycin introduction | Jul 2006<br>(95% CI Jan 2006 - Nov 2006) | Aug 1999<br>(95% CI Aug 1991 - Oct 2003) | Oct 1989<br>(95% CI Oct 1971 - Dec 1998) |
| **Aggregated log-likelihood** | -45.16 | -47.13 | -47.36 |
| **Aggregated AIC** | 94.32 | 98.26 | 98.72 |

AIC, Akaike information criterion; CI, confidence interval

## References

[1] Salado-Rasmussen K, Jensen JS. Mycoplasma genitalium Testing Pattern and Macrolide Resistance: A Danish Nationwide Retrospective Survey. Clin Infect Dis 2014; 59:24–30.

[2] Pond MJ, Nori A V, Witney AA, Lopeman RC, Butcher PD, Sadiq ST. High prevalence of antibiotic-resistant mycoplasma genitalium in nongonococcal urethritis: The need for routine testing and the inadequacy of current treatment options. Clin Infect Dis 2014; 58:631–7.

[3] Murray GL, Bradshaw CS, Bissessor M, Danielewski J, Garland SM, Jensen JS, et al. Increasing Macrolide and Fluoroquinolone Resistance in Mycoplasma genitalium. Emerg Infect Dis 2017; 23:809–12.

[4] Gesink DC, Mulvad G, Montgomery-Andersen R, Poppel U, Montgomery-Andersen S, Binzer A, et al. Mycoplasma genitalium presence, resistance and epidemiology in Greenland. Int J Circumpolar Health 2012; 71:1–8.

[5] Taylor-Robinson D, Jensen JS. Mycoplasma genitalium: from Chrysalis to Multicolored Butterfly. Clin Microbiol Rev 2011; 24:498–514.

[6] Wiesenfeld HC, Manhart LE. Mycoplasma genitalium in Women: Current Knowledge and Research Priorities for This Recently Emerged Pathogen. J Infect Dis 2017; 216:S389–95.

[7] Gaydos CA. Mycoplasma genitalium: Accurate Diagnosis Is Necessary for Adequate Treatment. J Infect Dis 2017; 216:S406–11.

[8] Bradshaw CS, Jensen JS, Waites KB. New Horizons in Mycoplasma genitalium Treatment. J Infect Dis 2017; 216:S412–9.

[9] Jensen JS, Bradshaw CS, Tabrizi SN, Fairley CK, Hamasuna R. Azithromycin Treatment Failure in Mycoplasma genitalium–Positive Patients with Nongonococcal Urethritis Is Associated with Induced Macrolide Resistance. Clin Infect Dis 2008; 47:1546–53.

[10] Bissessor M, Tabrizi SN, Twin J, Abdo H, Fairley CK, Chen MY, et al. Macrolide Resistance and Azithromycin Failure in a Mycoplasma genitalium-Infected Cohort and Response of Azithromycin Failures to Alternative Antibiotic Regimens. Clin Infect Dis 2015; 60:1228–36.

[11] Anagrius C, Loré B, Jensen JS. Treatment of Mycoplasma genitalium. Observations from a Swedish STD Clinic. PLoS One 2013; 8:e61481.

[12] Ito S, Shimada Y, Yamaguchi Y, Yasuda M, Yokoi S, Ito SI, et al. Selection of Mycoplasma genitalium strains harbouring macrolide resistance-associated 23S rRNA mutations by treatment with a single 1 g dose of azithromycin. Sex Transm Infect 2011; 87:412–4.

[13] Twin J, Jensen JS, Bradshaw CS, Garland SM, Fairley CK, Min LY, et al. Transmission and Selection of Macrolide Resistant Mycoplasma genitalium Infections Detected by Rapid High Resolution Melt Analysis. PLoS One 2012; 7:e35593.

[14] Falk L, Enger M, Jensen JS. Time to eradication of Mycoplasma genitalium after antibiotic treatment in men and women. J Antimicrob Chemother 2015; 70:3134–40.

[15] Read TRH, Fairley CK, Tabrizi SN, Bissessor M, Vodstrcil L, Chow EPF, et al. Azithromycin 1.5g Over 5 Days Compared to 1g Single Dose in Urethral Mycoplasma genitalium: Impact on Treatment Outcome and Resistance. Clin Infect Dis 2017; 64:250–6.

[16] Martin DH, Manhart LE, Workowski KA. Mycoplasma genitalium From Basic Science to Public Health: Summary of the Results From a National Institute of Allergy and Infectious Disesases Technical Consultation and Consensus Recommendations for Future Research Priorities. J Infect Dis 2017; 216:S427–30.

[17] Bonhoeffer S, Lipsitch M, Levin BR. Evaluating treatment protocols to prevent antibiotic resistance. PNAS Proc Natl Acad Sci United States Am 1997; 94:12106–11.

[18] Fingerhuth SM, Bonhoeffer S, Low N, Althaus CL. Antibiotic-Resistant Neisseria gonorrhoeae Spread Faster with More Treatment, Not More Sexual Partners. PLoS Pathog 2016; 12:1–15.

[19] Smieszek T, White PJ. Apparently-Different Clearance Rates from Cohort Studies of Mycoplasma genitalium Are Consistent after Accounting for Incidence of Infection, Recurrent Infection, and Study Design. PLoS One 2016; 11:1–18.

[20] Baumann L, Cina M, Egli-Gany D, Goutaki M, Halbeisen FS, Lohrer G-R, et al. Prevalence of Mycoplasma genitalium in different population groups: systematic review and meta-analysis. Sex Transm Infect 2018; 94:255–62.

[21] Horner P, Ingle SM, Garrett F, Blee K, Kong F, Muir P, et al. Which azithromycin regimen should be used for treating Mycoplasma genitalium? A meta-analysis. Sex Transm Infect 2018; 94:14–20.

[22] Miller JJ. The Inverse of the Freeman-Tukey Double Arcsine Transformation. Am Stat 1978; 32:138.

[23] Barendregt JJ, Doi SA, Lee YY, Norman RE, Vos T. Meta-analysis of prevalence. J Epidemiol Community Health 2013; 67:974–8.

[24] Akaike H. A New Look at the Statistical Model Identification. IEEE Trans Automat Contr 1974; 19:716–23.

[25] Chrisment D, Charron A, Cazanave C, Pereyre S, Bébéar C. Detection of macrolide resistance in Mycoplasma genitalium in France. J Antimicrob Chemother 2012; 67:2598–601.

[26] Touati A, Peuchant O, Jensen JS, Bébéar C, Pereyrea S. Direct Detection of Macrolide Resistance in Mycoplasma genitalium Isolates from Clinical Specimens from France by Use of Real-Time PCR and Melting Curve Analysis. J Clin Microbiol 2014; 52:1549–55.

[27] Le Roy C, Hénin N, Pereyre S, Bébéar C. Fluoroquinolone-Resistant Mycoplasma genitalium, Southwestern France. Emerg Infect Dis 2016; 22:1677–9.

[28] Joly-Guillou M-L, Lasry S. Practical Recommendations for the Drug Treatment of Bacterial Infections of the Male Genital Tract Including Urethritis, Epididymitis and Prostatitis. Drugs 1999; 57:743–50.

[29] Björnelius E, Magnusson C, Jensen JS. Mycoplasma genitalium macrolide resistance in Stockholm, Sweden. Sex Transm Infect 2016; 93:167–8.

[30] Birger R, Saunders J, Estcourt C, Sutton AJ, Mercer CH, Roberts T, et al. Should we screen for the sexually-transmitted infection Mycoplasma genitalium? Evidence synthesis using a transmission-dynamic model. Sci Rep 2017; 7:16162.

[31] Unemo M, Shafer WM. Antimicrobial resistance in Neisseria gonorrhoeae in the 21st Century: Past, evolution, and future. Clin Microbiol Rev 2014; 27:587–613.

[32] Chow JW, Fine MJ, Shlaes DM, Quinn JP, Hooper DC, Johnson MP, et al. Enterobacter Bacteremia: Clinical Features and Emergence of Antibiotic Resistance during Therapy. Ann Intern Med 1991; 115:585–90.

[33] Carmeli Y, Troillet N, Eliopoulos GM, Samore MH. Emergence of Antibiotic-Resistant Pseudomonas aeruginosa?: Comparison of Risks Associated with Different Antipseudomonal Agents. Antimicrob Agents Chemother 1999; 43:1379–82.

[34] Unemo M, Jensen JS. Antimicrobial-resistant sexually transmitted infections: gonorrhoea and Mycoplasma genitalium. Nat Rev Urol 2017; 14:139–52.

[35] Golden MR, Workowski KA, Bolan G. Developing a Public Health Response to Mycoplasma genitalium. J Infect Dis 2017; 216:S420–6.

[36] Read TRH, Fairley CK, Murray GL, Jensen JS, Danielewski J, Worthington K, et al. Outcomes of Resistance-guided Sequential Treatment of Mycoplasma genitalium Infections: A Prospective Evaluation. Clin Infect Dis 2018; doi:10.1093/cid/ciy477.

